# Fast Probabilistic Whitening Transformation for Ultra-High Dimensional Genetic Data

**DOI:** 10.1101/2025.09.01.673591

**Authors:** Gabriel E. Hoffman, Christian P. Dillard, Kiran Girdhar, Panos Roussos

## Abstract

Statistical methods often make assumptions about independence between the samples or features of a dataset. Yet correlation structure is ubiquitous in real data, so these assumptions are often not met in practice. Whitening transformations are widely applied to remove this correlation structure. Existing approaches to whitening are based on standard linear algebra, rather than a probabilistic model, and application to high dimensional datasets with *n* samples and *p* features is problematic as *p* approaches or exceeds *n*. Moreover, the computational time becomes prohibitive since the naive transform is cubic in *p*. Here we propose a probabilistic model for data whitening and examine its properties based on first principles as *p* increases. We demonstrate the statistical properties of the probabilistic model and derive a remarkably efficient algorithm that is linear instead of cubic time in the number of features. We examine the out-of-sample performance of the probabilistic whitening model on simulated data, and real genotype data. In an application to impute z-statistics from unobserved genetic variants from a genome-wide association study of schizophrenia, the probabilistic whitening transformation, had the lowest mean square error while being up to an order of magnitude faster than other methods. Using this approach, we also identify tandem repeats that explain genetic regulatory signals for disease-relevant genes. Analyses are implemented in our novel open source R packages `decorrelate` and `imputez`.

## 1 Introduction

Correlation structure is ubiquitous in real world datasets. Yet statistical models often make assumptions about independence between a dataset’s rows or columns that are not satisfied practice. Whitening transformations are a widely used preprocessing step to remove correlation structure in the data before performing other analyses [1]. In general, for a data matrix *Y*_*n×p*_ with *n* samples and *p* features, the procedure computes the sample covariance matrix between features, Σ_*p×p*_, and transforms the observed data by post-multiplying by the non-unique whitening matrix 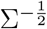 defined using standard linear algebra [1]. Transforming rows instead follows a complementary procedure. Covariance and their corresponding whitening matrices, along with correlation and their corresponding ‘decorrelating’ matrices, play a central role in many standard analysis methods such as multivariate regression, canonical correlation analysis, linear and quadratic discriminate analysis, Gaussian mixture models, Maholanobis distance, multivariate outlier detection, and independent component analysis [2–4], linear mixed models [5, 6], as well as deep learning applications [7–9].

For low dimensional data where *n* ≫ *p* seen in classical statistical settings, whitening transforms the data to have an identity sample covariance matrix. This procedure assumes that the true covariance matrix is known, or is well estimated by the sample covariance matrix. Yet the use of the sample covariance matrix for this transformation can be problematic for two distinct reasons. Computationally, evaluating the covariance matrix and creating the whitening matrix requires *O* (*p*^2^) memory and *O* (*p*^3^) time, which becomes infeasible for large *p* [1]. Statistically, the transformation must be adapted for the high dimensional case since the sample covariance matrix is no longer full rank or invertible. In modern high dimensional datasets, these computational and statistical limitations raise issues for the application of whitening transforms either as a preprocessing step, or a central part of widely used statistical methods.

Beyond general applications in statistical analysis, the field of statistical genetics makes extensive use of whitening transformations in order to account for the correlation between nearby genetic variants. This correlation structure is termed ‘linkage disequlibirum’ or ‘LD’ and is a major complication in downstream analysis of regression summary statistics (i.e. coefficient estimates or z-statistics) from genome-wide association studies [10]. It is common practice to estimate the correlation matrices from an external ‘reference panel’ of samples not included in the study of interest [10]. These ‘out-of-sample’ estimates are then used either as a preprocessing step to transform the summary statistics to be approximately uncorrelated, or internally in a statistical model. This approach has become standard in the field for phenotype prediction [11, 12], statistical fine-mapping [13–15], imputing z-statistics [16], detecting anomalous z-statistics [17], multiple testing correction [18], identifying genome annotations enriched for risk variants [19], and simulating knockoff datasets with similar correlation structure [20]. While the literature on whitening transformations has focused on estimating and applying the transform on the same data [1], an understanding of the performance of these out-of-sample applications has been lacking [21].

Direct estimation of covariance and precision matrices is a huge field of research [22–24]. Many approaches apply regularization to ensure the estimated covariance or precision matrix has specific properties. Widely used methods apply regularization so that the estimated covariance matrix is positive definite by shrinking the sample eigen-values [25, 26], or ensure sparsity of the covariance matrix and its inverse by shrinking regression coefficients corresponding to partial correlation coefficients [27]. Others regularize the condition number [28]. There have been many methods formulated as a convex combination of two matrices. Some methods use a convex combination between a non-sparse and sparse components [29, 30]. Stein-type estimators use a convex combination between the sample covariance and a target matrix [25, 26, 31]. A probabilistic approach modeling the observed data as a multivariate Gaussian with an inverse Wishart prior on the covariance gives a posterior mean that is a convex combination between the sample covariance matrix and the prior scale matrix [32–36].

Yet whitening transformations used in a preprocessing step or internally in a statistical model do not require *direct* estimates of the covariance matrix. Instead, covariance matrices are merely nuisance parameters: essential to the model, but not of direct interest. In data whitening, only the transformed data, rather the covariance matrix itself, is of interest. Here we seek to account for the covariance, rather than estimate it directly, and we propose an implicit covariance approach. This important distinction motivates the probabilistic framework, statistical theory and computationally efficient algorithms we use here to perform whitening transformations without directly estimating the covariance or whitening matrices.

Bayesian or, more broadly, probabilistic formulations of existing techniques can lead to deeper understanding of model behavior, generalization of the model, regularization to incorporate prior knowledge, or incorporation into a hierarchical model [3, 4]. Widely used Gaussian mixture models and extensions follow from a probabilistic formulation of the k-means problem, and probabilistic formulations of principal components analysis have long been influential in statistical machine learning [37, 38]. Yet we are not aware of such a rigorous probabilistic formulation for the widely used whitening transformation.

Here we propose a probabilistic formulation of the whitening transformation under Gaussian noise, and incorporate regularization in the form of an inverse-Wishart prior distribution on the covariance matrix. This produces a well-known estimator with a single hyperparameter that can be efficiently estimated using an empirical Bayes approach. Using first principles of the probabilistic model, we give a direct interpretation of the hyperparameter and examine the statistical behavior of this whitening transformation. We demonstrate that the probabilistic whitening transformation does not require direct estimation of the covariance or whitening matrices, and derive a remarkably efficient approach to apply the transformation to ultra-high dimensional data.

We examine the empirical performance of the probabilistic whitening transformation on simulated data and genetic data from the 1000 Genomes Project [40]. We apply the whitening transformation to a genome-wide association study of schizophrenia [41], and impute z-statistics of unobserved variants using an external reference panel. Finally, we use this approach to impute genetic regulatory signals for tandem repeats effecting gene expression in the human brain.

## 2 Results

### 2.1 Introduction to whitening transformations

Consider observed data *Y*_*n×p*_ with *n* samples and *p* features drawn from a distribution with mean vector 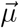, identity covariance between rows, and covariance between features given by Σ_*p×p*_. Let

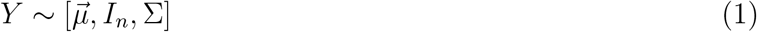

indicate these properties without specifying particular distributional assumptions. Transforming the data by the non-unique whitening matrix 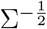 produces mean and covariances according to

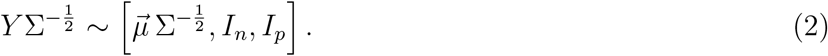

where Σ is known. This follows from the definition of covariance and the standard property that 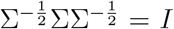. While it is common to assume a multivariate Gaussian distribution for the observed data, this transformation applies to the first two moments of any multivariate distribution. Applying this transformation scales and rotates the data to produce an identity covariance matrix. The whitening matrix is not uniquely defined and different transformation matrices result in different rotations all with identity covariance [1].

The sample covariance is 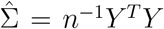, following mean centering of the columns. Let the singular value decomposition of the scaled observed data be

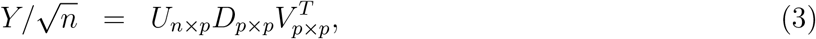

with singular values 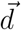, so that the eigen decomposition of the sample covariance is 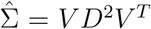 and the whitening matrix 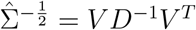 transforms the data to have identity covariance (**SI 5.1**).

This definition of the whitening matrix minimizes the total squared distance between original and transformed data, and is termed the ‘zero-phase components analysis’ or ZCA whitening [1]. This three step transform, one for each matrix multiplication, can be interpreted as rotating, scaling, then unrotating the observed data to give an identity sample covariance (**Figure 1**). This whitening transform is the most intuitive since it returns the data to its original axes, and it is the foundation for the transformations we describe below.

**Figure 1.**
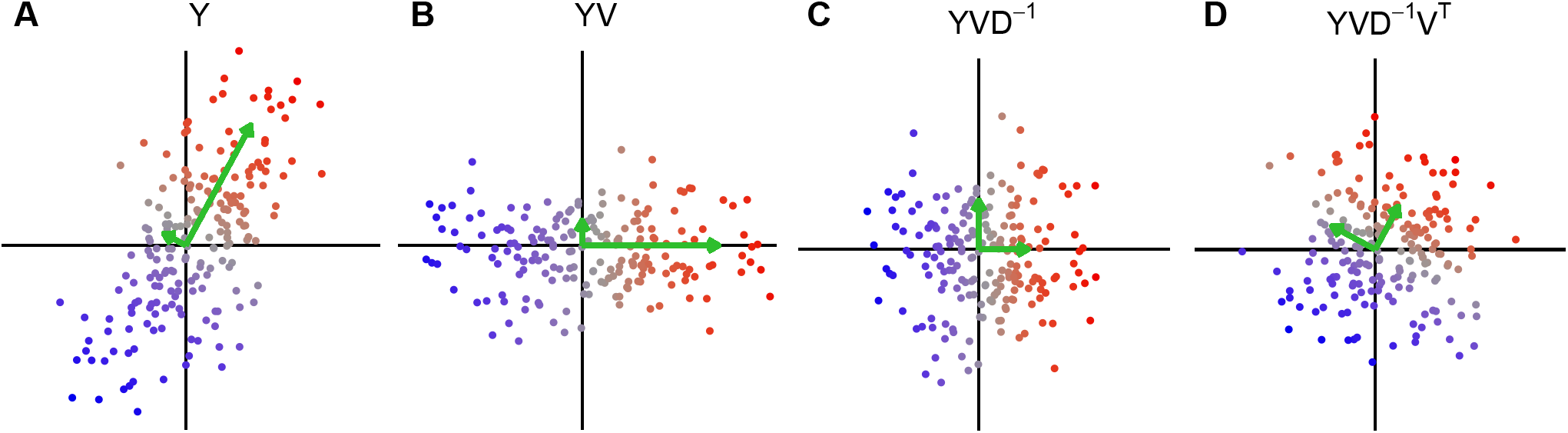
Intuition for data whitening transformation. **A)** Original data, **B)** Data rotated along principal components, **C)** Data rotated and scaled, **D)** Data rotated, scaled and rotated back to original axes. Green arrows indicate principal axes and lengths indicate eigen-values.

In practice, the true value of Σ is not known and it must be estimated from the observed data *Y*. An exact whitening transformation that produces an identity covariance matrix is only possible when *n > p* so that that sample covariance matrix, 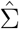, is invertible. Even in this case, the transform can be unstable when the trailing singular-values are very small, since evaluating 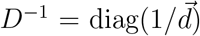 becomes very unstable for small values of *d*_*i*_ and are sensitive to precision limitations in numerical linear algebra. In high dimensional data with *n < p*, the trailing singular values are exactly zero, so some form of regularization is needed. An exact whitening transformation no longer exists in these cases, we want to learn a transformation that that transformed data has a covariance close to identify by some metric (**SI 5.2**).

### 2.2 A probabilistic model for whitening transformations

Here we propose a whitening transform based on an probabilistic model of the observed data and demonstrate superior statistical performance compared to standard data whitening transformations [1], even in the low-dimension case where both are applicable. In proposing a probabilistic model, we consider the observed data to have a matrix normal distribution^1^ with unknown mean, identity covariance between rows, and unknown covariance between columns. We then place an inverse Wishart (IW) prior on the covariance matrix between columns. The normal assumption is of course natural for unbounded continuous data, and the IW is natural because it is the conjugate prior so that the posterior distribution of the covariance is also IW [42]. Formally,

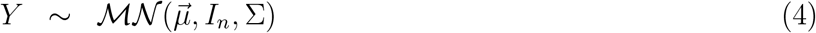

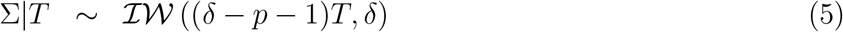

where *T* is the target matrix of the prior and *δ > p* − 1. Based on this model, the posterior expectation of the covariance is

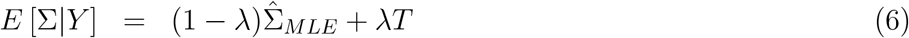

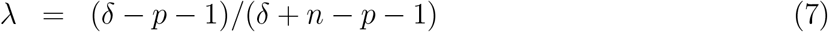

Where 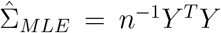 is the sample covariance matrix and *λ* ∈ [0, 1] by definition [32–35] The posterior expectation is therefore a convex combination between the sample covariance, 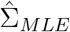, and the prior target matrix, *T*.

Setting the target matrix to be symmetric positive definite (SPD) ensures that the posterior expected covariance is also SPD [33]. Restricting *T* to be a minimally informative prior target corresponds to setting all prior covariances to zero. We focus here on the important case where the IW prior is centered around a scaled identity matrix, and we demonstrate that the estimate has exceptionally efficient computational scaling and produces estimates that are tractable to study analytically. Following previous work [25, 26, 33–35, 43] we set the target matrix to be *T* = *νI* where 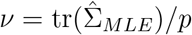 is the average variance across all *p* features. This gives the important special case we consider here with

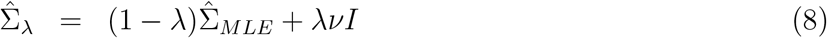

where 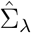 is defined as the posterior expected covariance from the probabilistic model.

In the important case of out-of-sample whitening where the whitening transform is learned from a training set before being applied to the observed data, the standard whitening transform motived by linear algebra assumes the correlation structure of the training and observed data are the same. This is a strong assumption that is rarely satisfied in practice. The finite size of the training set means that correlation structure will differ from the observed data, so the standard whitening transform will overfit the training data. Instead, the probabilistic model substantially relaxes this central assumption, and assumes only that the training and observed data are drawn from the same distribution.

In practice, the shrinkage can be applied to the correlation matrix while retaining the diagonal variance terms (**SI 5.3**).

### 2.3 Estimating the shrinkage parameter

Covariance estimators that are convex combinations of the sample covariance and target matrices have received much attention. The estimator for *λ* depends on assumptions about the data and the specified loss function. Ledoit and Wolf [25] and Schäfer and Strimmer [26] propose minimizing the expected squared Frobenius loss between the true and estimated covariance matrices, then use a consistent estimate of *λ* computed from the observed data. Further work extended this approach to improve estimation of *λ* in the high dimensional setting [31, 43, 44]. Others use a probabilistic model [32–35], cross-validation [45, 46] or the out-of-sample likelihood [47, 48]. Yet these methods are designed for direct estimation of covariance or precision matrices, rather than data whitening.

Following the probabilistic formulation, we take a Bayesian approach to estimating *λ*, and we refer to this as the Gaussian Inverse Wishart empirical Bayes (GIW-EB) model. We integrate out the covariance matrix and maximize the likelihood of the observed data as a function of *λ*. Due to the analytic properties of the Gaussian and IW distributions, the covariance matrix can be treated as a nuisance parameter and marginalized out according to

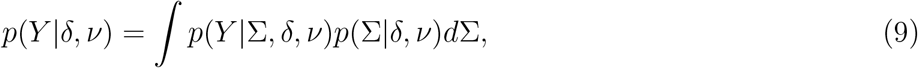

so the marginal distribution is multivariate Student-t [42]. The hyperparameter *δ*, and therefore *λ*, can be estimated directly from data using an empirical Bayes approach by maximizing *p*(*Y* | *δ, ν*) (**SI 5.4**). Given the special form of the target matrix, *νI*, we demonstrate that the data only enters the log-likelihood through the sample singular values. This has important implications for the probabilistic whitening transformation described below.

### 2.4 Properties of posterior expected covariance

The behavior of this probabilistic whitening transformation corresponding to the posterior expected covariance in Equation (8) can be understood in terms of the spectral decomposition of the observed data matrix, *Y*.

Based on the singular value decomposition of the observed data, the eigen decomposition of 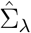 is

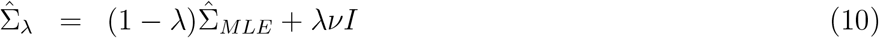

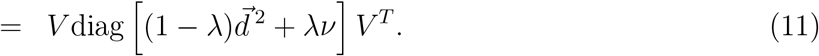

Therefore the eigen vectors of 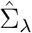 are the same as for the sample covariance matrix, but the eigen-values are shrunk so that the *j*^*th*^ eigen-value is 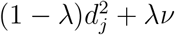. Since *λ* ∈ [0, 1], *ν >* 0, and 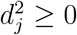, it follows that the eigen-values are always positive, so that 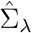 is aways positive definite and invertible, even when 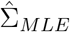 is not.

### 2.5 Statistical properties of probabilistic whitening transformation

The probabilistic whitening transformation can be understood in similar terms as the posterior expected covariance. Consider a fixed (i.e. non-random) latent data matrix *X*_*n×p*_ with uncorrelated columns, and let the observed data *Y*_*n×p*_ be generated by applying a rotation based on a symmetric positive definite covariance matrix Σ_*p×p*_ so that

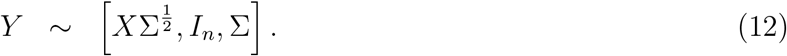

Given the observed data, *Y*, estimate the covariance matrix as 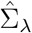 and apply the whitening transformation so that

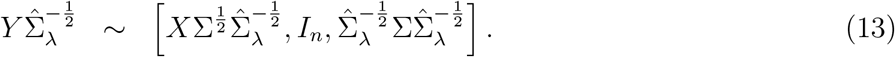

Since 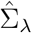 converges to Σ as *n* → ∞ for a fixed *p* [33], the whitening transform recovers the original signal according to

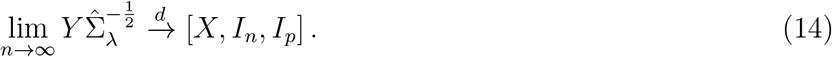

The whitening transformation can then be expressed in terms of singular vectors and modified singular values according to

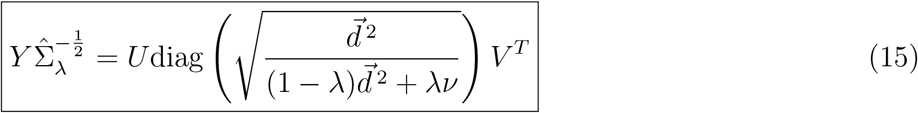

The central statement in Equation (15) of the whitening transformation in terms of the original singular vectors and modified singular values has both important statistical and computational implications we discuss below.

The whitening transformation retains the original singular vectors and acts only by modifying the sample singular values. From here we can examine the behavior of the transformation as a function of *λ* (**SI 5.5**), and interpret the effect of *λ* on the effective number of parameters used in covariance estimation (**SI 5.6**).

### 2.6 Scalar summaries of the correlation or covariance matrix

Common scalar summary statistics of covariance matrices can be computed efficiently based on the form of the spectral decomposition of 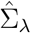 (**Table 1**). Standard values like trace, log determinant and condition number can be computed easily from the shrunken eigen-values. In addition the effective degrees of freedom [49], average correlation, average squared correlation [50], effective variance [51], effective number of features [52] are easily computed (**SI 5.7**).

**Table 1.** Computing scalar summary statistics of correlation matrices. Terms are defined in the text except for letting the shrunken eigen-values be 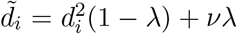, and the mean shrunken eigen-value be 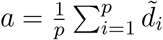. Let 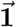 be a vector of length *p* with all entries having value 1.

| Statistic | Formula | Reference |
| --- | --- | --- |
| Trace | $\sum_{i=1}^p \tilde{d}_i$ | |
| Log det | $\sum_{i=1}^p \log \tilde{d}_i$ | |
| Condition number | $\tilde{d}_1 / \tilde{d}_p$ | |
| Average squared correlation | $\sum_{i=1}^p (\tilde{d}_i - a)^2 / p(p-1)a^2$ | [50] |
| Effective degrees of freedom | $(1 - \lambda)p(p-1)/2$ | [49] |
| Average correlation | $(\vec{1}^T \Sigma_\lambda \vec{1} - p) / (2p(p-1))$ | |
| Effective variance | $\exp \left[ \frac{1}{p} \sum_{i=1}^p \log \tilde{d}_i \right]$ | [51] |
| Effective number of features | $\vec{1}^T \Sigma_\lambda^{-1} \vec{1}$ | [52] |

### 2.7 Computational methods for data whitening

The computational time for applying the standard whiting transformation, even in the low dimensional setting, can quickly become intractable for moderate *p*. The time complexity of the standard algorithm is *O* (*p*^3^ + *np*^2^) (**SI 5.8**), and becomes very expensive for *p >* 100 and can be intractable for *p >* 2000 features. Yet the entire procedure can be substantially accelerated using properties linear algebra

(**Table 2**). It is not necessary to explicitly compute the sample covariance and whitening matrices and then transform the observed data. Equation (15) rephrases the whitened data in terms of modified singular values, and the original left and right singular vectors. This gives a substantially faster algorithm. In the low dimensional setting of *n > p*, computing the SVD of the observed data is *O*(*np*^2^), estimating and applying the *ν* and *λ* is negligable, and evaluating the matrix products is *O* (*np*^2^). The procedure is thus substantially reduced to *O* (*np*^2^) and uses *O* (*np*) memory.

**Table 2.** Computational and memory complexity of whitening methods for *n* samples and *p* features using a rank *k* transformation.

| Method | Decomposition | Estimate $\lambda$ | Transform | Total | Memory |
| --- | --- | --- | --- | --- | --- |
| Standard | $p^3 + np^2$ | $\geq np$ | $np^2$ | $p^3 + np^2$ | $p^2$ |
| GIW-EB ($n > p$) | $np^2$ | $n$ | $np^2$ | $np^2$ | $np$ |
| GIW-EB ($n < p$) | $n^2p$ | $p$ | $n^2p$ | $n^2p$ | $np$ |
| GIW-EB, low rank | $knp$ | $k$ | $knp$ | $knp$ | $np$ |

#### 2.7.1 Accelerating computation in the low dimensional setting

#### 2.7.2 Accelerating computation in the high dimensional setting

The standard formulation of the data whitening transformation does not apply to the high dimensional setting of *n < p* since the sample covariance matrix is no longer invertible. Some regularization of the singular values is necessary. The probabilistic whitening transformation indeed shrinks the singular values towards *ν* and addresses the statistical challenge of the high dimensional setting. Yet this computational approach is still quadratic in *p*, and is infeasible for large *p* typical in real datasets.

In general, the rank, *r*, of the sample covariance matrix is bounded above by min(*n, p*). In the high dimensional setting, at most only *r* ≤ *n* singular values are nonzero. Evaluating the SVD is *O* (*n*^2^*p*) in this case only produces the first *n* singular vectors and values. So there remain *p* − *n* uncomputed singular vectors with corresponding singular values of zero. Directly computing these is impractical and, in fact, unnecessary.

Again, the key insight comes from Equation (15). Since all *p* singular values, including ones with value zero, are modified to be positive it seems that even left and right singular vectors with no variation (i.e. zero singular values) are required to compute the transformation. Yet here we demonstrate that in fact only non-zero singular values and their corresponding singular vectors are required. This insight further reduces the computational complexity of the entire transform.

The substantial improvement in computational complexity for the whitening transformation in the high dimensional settings comes from computing and using only singular vectors corresponding to nonzero singular values. The method is useful beyond this specific context, so here we derive a general theorem and discuss special cases and applications. The theorem is exact if all nonzero singular values are retained, and is approximate of only a partial set are retained. The derivation generalizes and simplifies the work of Lippert, et al. [6] (**SI 5.9**).

**Theorem 1.** *Let C be a p* × *p symmetric positive semi-definite matrix, with k* ≤ *p positive eigen values. Let the eigen decomposition of C be* 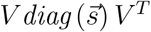 *where s*_*i*_ = 0 *for i > k. Let the eigen-vectors be partitioned as V* = [*V*_1_ *V*_2_], *and the eigen-values be partitioned as* 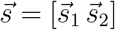 *to separate the nonzero and zero components. Let raising a matrix to an exponent α correspond to raising the eigen-values of the matrix to α. Then*

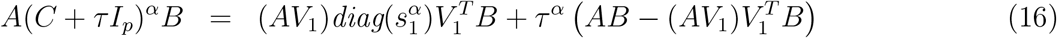

*where V*_1_ *and* 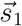 *indicate the first k eigen-vectors and values, respectively, and A*_*a×p*_ *and B*_*p×b*_ *are arbitrary matrices, τ is a positive scalar, and I*_*p*_ *is the p-dimensional identity matrix*.

Applying Theorem 1 to the whiting transform, and using the decomposition

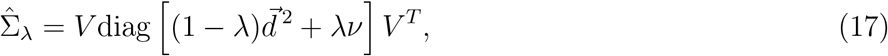

restated here from Equation (11), it follows that

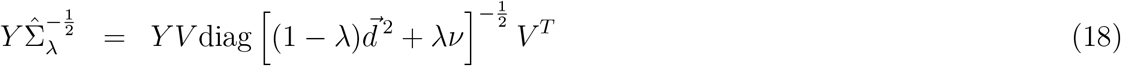

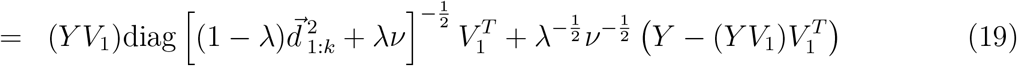

where *V*_1_ stores the first *k* eigen vectors of Σ_*λ*_ corresponding to the right singular vectors of 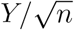 and 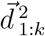 are the first *k* eigen values.

Using standard properties of matrix multiplication, the time complexity of evaluating these terms are defined as follows.

**Corollary 1.1**. *Since the time complexity of multiplying a* × *b and b* × *c matrices is O* (*abc*), *the time to evaluate the general formulation of Theorem 1 is O* (*abnp*), *once the SVD is already computed. In probabilistic data whitening in Equation (19), B* = *I and a* = *n so the time complexity reduces to O* (*knp*).

In general, the whitening transformation can be applied in *O* (*knp*) time after the SVD is computed. When the first *n* singular vectors and values are computed using the standard SVD, then *k* = *n*, so the entire procedure is *O* (*n*^2^*p*), which is now *linear* in the number of features. This is a dramatic improvement from the naive method that is *cubic* in the number of features (**Table 2**).

#### 2.7.3 Scaling to the ultra-high dimensional setting

While this algorithm is now linear in *p*, it is still quadratic in the number of samples, *n*. Yet empirical data can often be well approximated as a low rank matrix [53]. As *n* and *p* increase but the dimension of the underlying signal remains fixed, it is not necessary to compute all min(*n, p*) singular values and vectors. Instead a rank *k* ≪ min(*n, p*) partial SVD can be computed in *O* (*knp*) time to provide a rank *k* approximation of the observed data [54]. Moreover, the algorithm in Equation (19) can also be evaluated in *O* (*knp*) time given the partial SVD (Table 2). The complexity of this approach is now linear in both *n* and *p*, and dependent on the rank *k* approximation chosen by the analyst.

### 2.8 Beyond the whitening transformation

While this work focuses on the whitening transformation, applying Theorem 1 with other values for the general exponent *α* are also useful in practice. In the context of probabilistic whitening transformation with variables defined as in Equation (19), four common operations involving 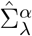 can be accelerated:

- *α* = 1 gives the posterior expected covariance matrix.
- *α* = −1 gives the corresponding precision matrix.
- 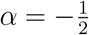 gives the whitening transformation.
- 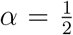 gives the ‘coloring matrix’ so that post-multiplying by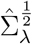 produces a correlation of 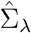 between columns.

## 3 Results

### 3.1 Performance on simulated data

The formulation of the probabilistic whitening transformation provides two advances of practical interest in analyses of large-scale datasets. First, the implicit covariance approach reduces computational time for high-dimensional data from the standard cubic time method that creates and then inverts the covariance matrix. For example, in simulations with *n* = 5000 and an increasing number of features the GIW-EB methods runs in < 2 min and the low rank GIW-EB with k=50 runs in < 1 min, when the standard method becomes intractable both in terms of time and memory (**Figure 2A**). Second, the probabilistic model motivates the empirical Bayes estimation of the shrinkage parameter which only uses the sample singular values of the data. Due to its probabilistic formulation, the whitening high-dimensional data using the GIW-EB method can produce lower out-of-sample error than other methods. For example, in a simulation Scenario 3 (described below), both full and low rank GIW-EB give estimated values of *λ* that approach the optimal value while other methods give *λ* values that are too small and have higher error (**Figure 2B**).

**Figure 2.**
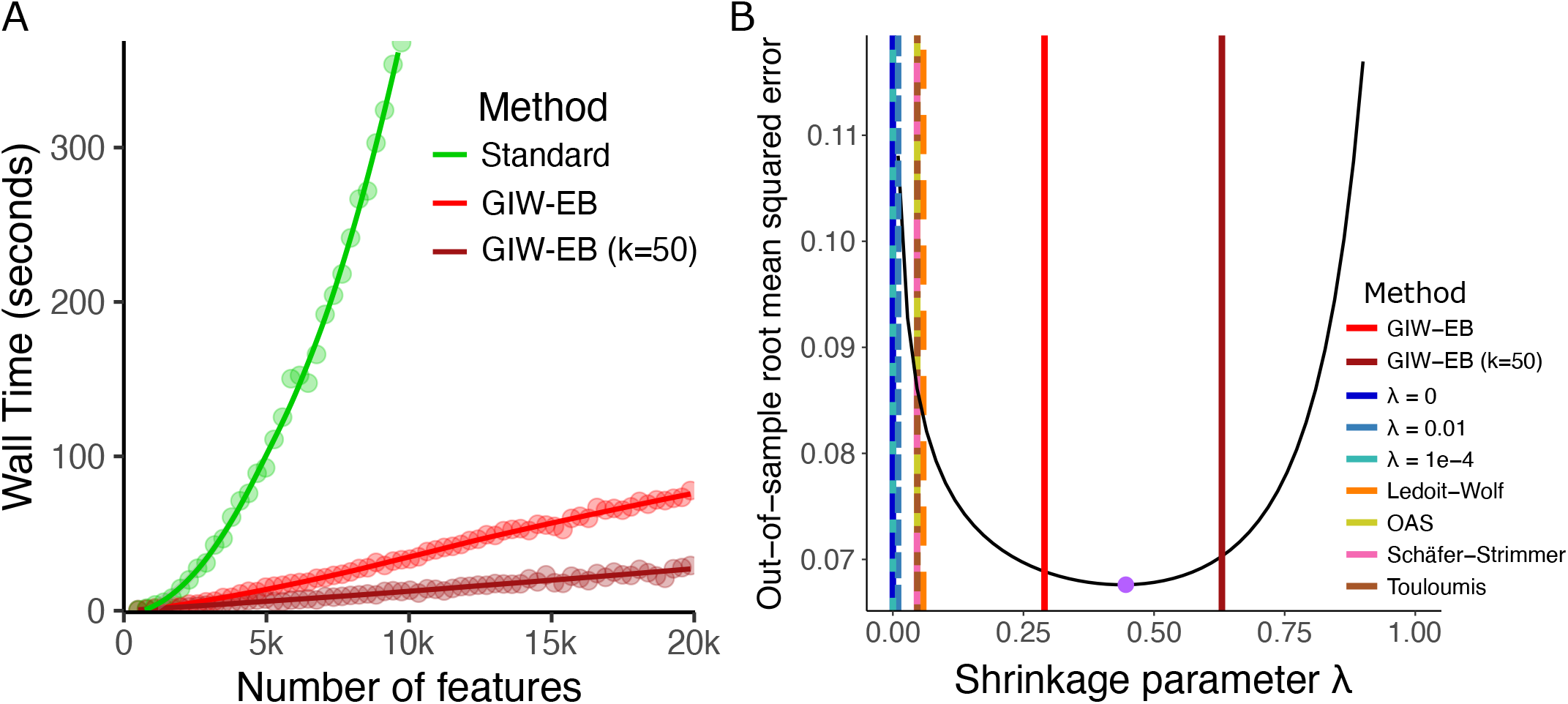
Performance of probabilistic whitening transformation. **A**) Run times are shown for standard (green), GIW-EB (red) and low rank GIW-EB with *k* = 50 (dark red) whitening transformations. Results shown for *n* = 5000 samples and an increasing number of features. Points indicate observed wall time and lines show loess smooth. **B**) The empirical out-of-sample root mean squared error for a simulated dataset across the range of *λ* values is shown by the black curve. The *λ* value giving the lowest error is show by the purple point. Values of *λ* estimated by other methods are shown by vertical lines.

Since an exact whitening transformation produces an identity covariance matrix when applied to the data it was estimated from, the accuracy of an approximate whitening transformation can be evaluated by estimating it from a subset of the data and computing the empirical covariance after applying the transform to the rest of the dataset. This out-of-sample accuracy of the whitening transformation is captured by the root mean squared error of the difference between the empirical covariance of the transformed data and the identity matrix.

We consider simulations with *n* = 500 and the number of features *p* increasing from 20 to 2000, with correlation between features in each of 4 scenarios:

1. constant correlation of *ρ* = 0.6
2. auto-correlation with *ρ* = 0.95
3. 4 blocks of constant correlation of *ρ* = 0.6
4. 4 blocks of auto-correlation with *ρ* = 0.95

In Scenario 3, most methods show equivalent performance for small values of *p*, error increases as *p* approaches *n*, and then error decreases and converges to the optimal root mean squared error (**Figure 3A**). The ‘Oracle’ method, shown in black here, transforms the data using the true covariance, so it gives a lower bound on the error. Using a whitening transformation without shrinkage by setting *λ* = 0 performs poorly even for small *p* and error only increases with *p*. Using the pseudoinverse improves performance when *p > n*, but does not compete with shrinkage methods. Examining different approaches to estimate the shrinkage parameter, the GIW-EB method gives the smallest error across all values of *p* and the low rank GIW-EB with k = 50 outperforms other methods for moderate *p*. Both full and low rank GIW-EB give errors that decrease with *p* and avoids the increase in error at *p* ≈ *n* seen with other shrinkage methods.

**Figure 3.**
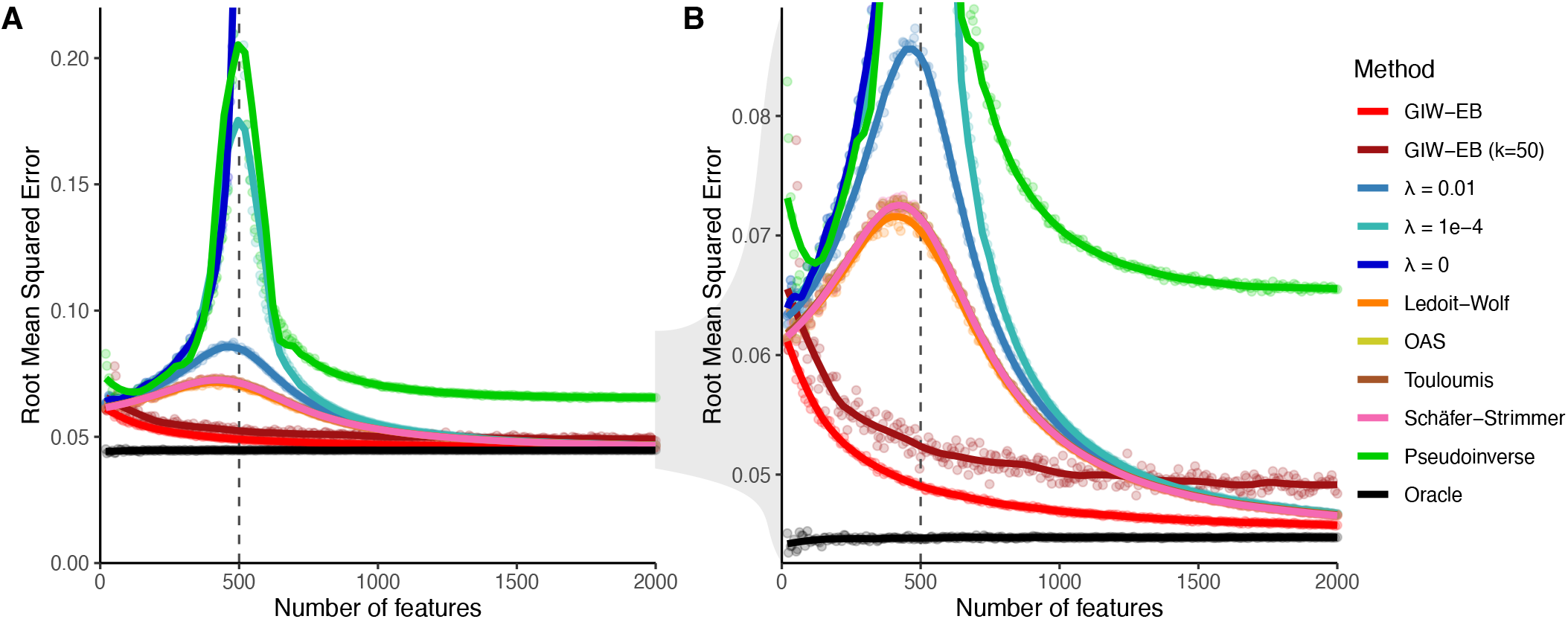
Out-of-sample performance of whitening transformation. Simulated data for *n* = 500 sample are drawn from a multivariate normal distribution with known covariance matrix and 11 whitening transformations are then applied to 500 additional samples from the same distribution. Of these, 9 methods either estimate the shrinkage parameter *λ* from the data or use a fixed value. Setting *λ* = 0 corresponds to no shrinkage, and pseudoinverse does not perform any shrinkage. The oracle method performs a whitening transformation using the true correlation matrix from which the data are simulated, and so gives the best out-of-sample performance. Performance is evaluated by comparing the correlation matrix of the transformed data to the identity matrix, and computing the root mean squared error (rMSE). **A**) rMSE of the whitening transformation for an increasing number of features **B**) Zoom-in on y-axis from (**A**). Points indicate observed values and lines show loess smooth.

Overall, the GIW-EB method consistently gives the lowest out-of-sample root mean square error across a range of *p* values in the 4 simulation scenarios (**Figure S1-4**). The low rank GIW-EB method is competitive in Scenarios 1 and 3, were the true covariance is low rank.

### 3.2 Performance on genetic data from 1000 Genomes Project

Since many analyses of summary statistics from genome-wide association studies apply a whitening transformation estimated from an external reference panel, we evaluated the out-of-sample performance of these whitening transformations on genetic data from the 1000 Genomes Project [40]. Individuals of European, Asian and African ancestry were analyzed separately, and the whitening transformation was performed on independent LD blocks including varying number of genetic variants defined in each population [55]. The whitening transform was trained on half of the samples and evaluated on the other half. Despite the genetic data not being multivariate normal, shrinkage methods performed well across most windows in European individuals (**Figure 4A**). The GIW-EB matches or is competitive with other methods that estimate *λ* from the data. The low rank GIW-EB method with *k* = 50 is most competitive for windows with *<* 1, 500 variants. The GIW-EB methods and methods with fixed *λ* values use the implicit covariance algorithm, and these give the best computational performance with a wall time of ∼ 5 min compared to 3-30 hours for the other methods (**Figure 4B**). Error profiles and computational time was similar for individuals of African and Asian ancestry (**Figure S5**).

**Figure 4.**
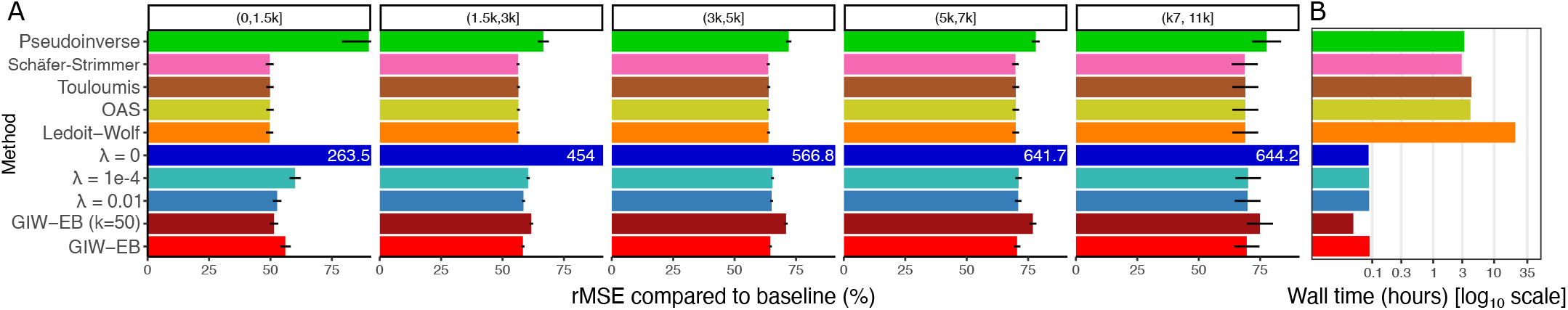
Out-of-sample whitening performance on genetic data. **A**) Root mean squared error (rMSE) compared to baseline for European individuals stratified by the number of genetic variants in the LD window. Values indicate the values averaged across windows, and bars indicate the 95% confidence interval. When normalized rMSE values exceed the axes, the value is shown in white text. **B**) Wall time for each method shown on a log_10_ scale. Methods using GIW-EB or fixed *λ* values use the implicit covariance algorithm.

### 3.3 Application to imputing GWAS summary statistics

Genome-wide associations studies (GWAS) release summary statistics from their data corresponding to z-statistics from the set of genetic variants analyzed in the study. Since the number of genetic variants analyzed in the original study can vary widely, there has been interest in imputing z-statistics of additional variants observed in an external reference panel. Since the correlation between z-statistics of two genetic variants matches the linkage disequilibrium between the two variants, Pasaniuc, et al. [16] proposed a method to impute z-statistics from unobserved variants using the set of observed z-statistics and the correlation structure between variants in a genomic region. This is substantially faster and cheaper than performing ‘genotype imputation’ on the full dataset [56].

Imputing z-statistics from unobserved variants can be stated in terms of out-of-sample whitening (**SI 5.10**), and we evaluated the performance of different whitening methods on imputation accuracy. We used summary statistics for 5.6M variants from schizophrenia GWAS of 58K cases and 77K controls of European ancestry [41]. Z-statistics for 15M variants were imputed using a reference panel of 407 unrelated individuals of European ancestry from the 1000 Genomes Project [40] using a sliding window of 1 Mb and a flanking region of 250 kb across all autosomes. Accuracy was evaluated by comparing the imputed and observed z-statistic for 460K randomly selected variants with minor allele frequency ≥ 5%. GIW-EB gave high imputation accuracy across a broad range of z-statistic values (**Figure 5A**), and give smallest genome-wide root mean squared error (**Figure 5B**). Imputation error increases for small minor allele frequencies (MAF), and GIW-EB gives the smallest squared error across the range of MAF values (**Figure 5C**). Given 407 samples and a mean of ∼ 3100 variants in each genome window, the low rank GIW-EB with *k* = 50 was not a good approximation in this case. The GIW-EB methods and methods with fixed *λ* values use the implicit covariance algorithm, and these give the best computational performance and require < 1 hr for genome-wide analysis for this reference panel compared to 4-126 hours for other methods (**Figure 5D**). Overall, GIW-EB method is the most accurate and fastest approach to impute GWAS summary statistics.

**Figure 5.**
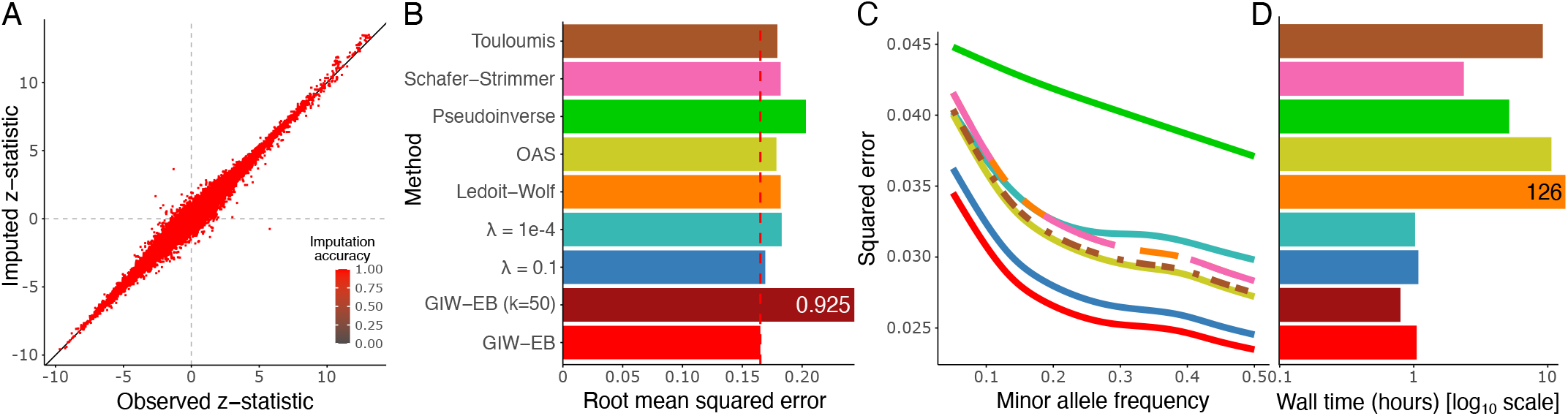
Imputation of GWAS summary statistics. **A**) Imputed z-statistics plotted against the observed z-statistics for 461,707 SNPs. **B**) Root mean squared prediction error for each method. Red vertical dashed line indicates the value for GIW-EB. **C**) Square prediction error as a function of minor allele frequency. Smooth curves are fit by a generalized additive model that is the default in `ggplot2::geom_smooth()`. GIW-EB (k=50) was excluded from this plot due to high imputation error. **D**) Compute time required for each method. When values exceed the axes, values are shown in text. Methods using GIW-EB or fixed *λ* values use the implicit covariance algorithm.

0.925

### 3.4 Application to impute eQTL signals for tandem repeats

Studies of genetic risk for disease and genetic regulation of gene expression have focused on common single nucleotide polymorphisms (SNPs) because they can be measured cheaply at scale. In recent work, Ziaei Jam, et al. [57] created a population reference panel including donors from 1000 Genomes Project of 1.7M tandem repeats that are not captured by existing panels. Here, we integrate this resource with summary statistics from a study of genetic regulation of gene expression in the human brain using almost 4K bulk RNA-seq samples [58]. As a proof of principle, we applied this approach to 3 disease-relevant genes with complex genome structure and we identified tandem repeats that explain the genetic regulatory signal (**Figure 6**). C9orf72 is a key risk gene for amyotrophic lateral sclerosis, with expansion of a intronic hexanucleotide repeat conferring disease risk and effecting gene expression[59]. TSPAN14 and TYK2 have been implicated in risk for Alzheimer’s disease [60], and here we identify tandem repeats with a stronger association with gene expression than SNPs in the region.

**Figure 6.**
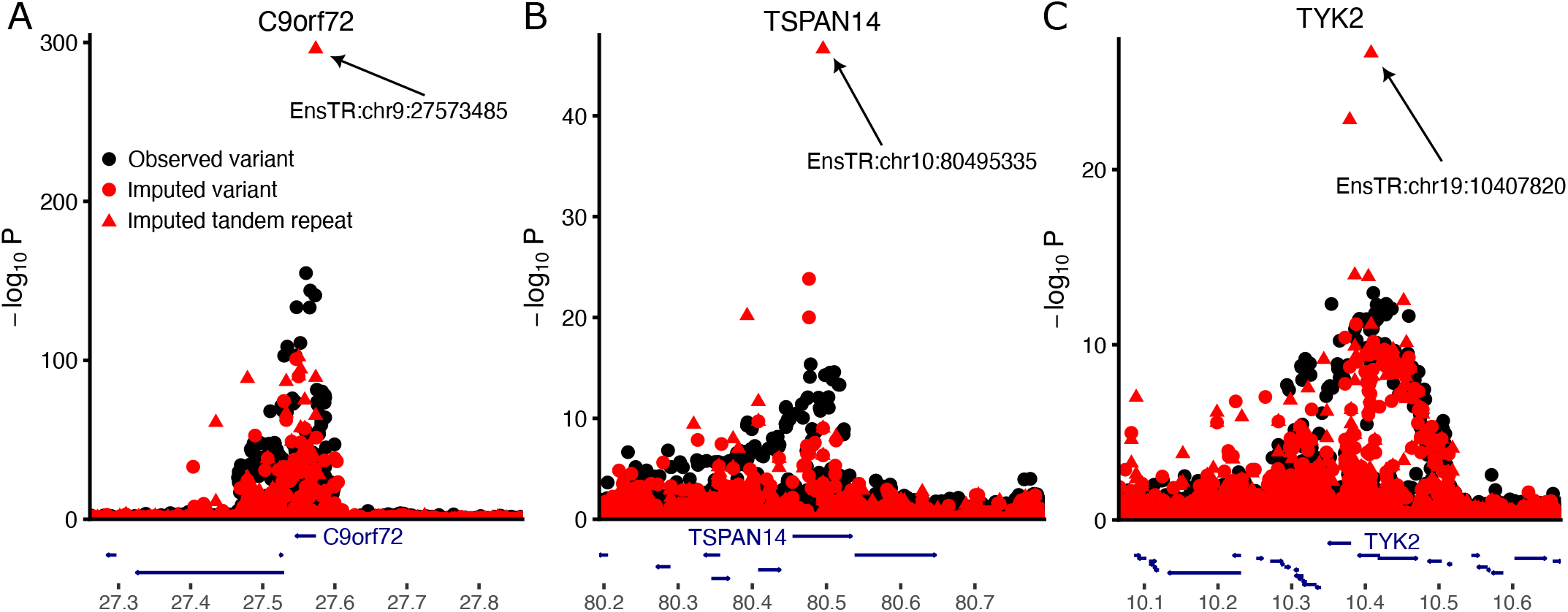
Imputation of eQTL summary statistics in human brain implicates tandem repeats‥. Results are shown for **A**) C9orf72, **B**) TSPAN14, and **C**) TYK2. Black points indicate observed variants included in the original eQTL analysis, red points indicates imputed variants, and red triangles indicate imputed tandem repeats.

## 4 Discussion

The whitening transformation is a central, if often unstated, component of many widely used statistical methods. While many standard statistical methods have been reformulated with a probabilistic model, or extended to handle regularization and modern high dimensional datasets, the whitening transform has received less attention. The standard whitening transformation is limited by computational time, memory usage, and gives poor out-of-sample performance on modestly sized datasets. Regularizing the covariance estimate improves statistical performance, but is still very computationally demanding for high dimensional datasets.

We proposed a regularized probabilistic whitening transformation and examined its statistical properties based on the singular value decomposition of the data matrix. The particular form of the probabilistic formulation enables an implicit covariance approach that avoids direct estimation of the covariance matrix and reduces the computational time from cubic to linear in the number of features. This can reduce computational time by a factor of ≥ 100 in practical applications.

In simulations and applications high-dimensional genotype data from the 1000 Genomes Project, we demonstrated that our GIW-EB method matches or exceeds other methods in out-of-sample whitening benchmarks. In an application to impute z-statistics from a schizophrenia GWAS, GIW-EB gave the best imputation performance. As a proof of principle, we demonstrate that z-statistic imputation powered by our GIW-EB whitening transform can identify tandem repeats explaining the genetic regulatory signals for disease-relevant genes for amyotrophic lateral sclerosis and Alzheimer’s disease. As the scale of genomic datasets continues to increase, the implicit covariance approach of the probabilistic model will enables application of whitening transforms on high dimensional data where existing methods are intractable. Future applications include single cell gene expression and larger genetic references panels.

## Analysis code

Code implementing the probabilistic whitening transformation with implicit covariance is available in the R package `decorrelate` from https://cran.r-project.org/package=decorrelate. Documentation is available at https://gabrielhoffman.github.io/decorrelate. Code implementing the imputation of summary statistics is available in the R package `imputez` available at https://gabrielhoffman.github.io/imputez. R packages have also been deposited at Zenodo: decorrelate https://doi.org/10.5281/zenodo.17426788, imputez https://doi.org/10.5281/zenodo.17426882. Code for simulation and analysis has been deposited at Zenodo https://doi.org/10.5281/zenodo.17427136 and https://doi.org/10.5281/zenodo.17427128. Times were evaluated using 12 cores on a 48 core Intel Xeon Platinum 8462Y CPU @ 2.8GHz running R v4.3.3 linked to a parallelized Intel Math Kernel Library for linear algebra operations.

## Acknowledgements

We thank Bernie Devlin for valuable feedback.

## 5 Supplementary Information

### 5.1 Defining the whitening matrix

Consider data matrix *Y*_*n×p*_ covariance between columns given by Σ_*p×p*_ and each column having zero mean after centering. Defining the singular value decomposition 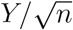 as *UDV* ^*T*^, then

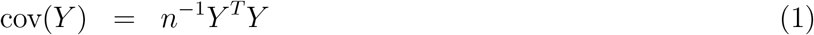

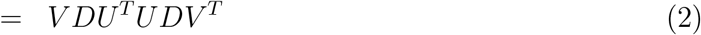

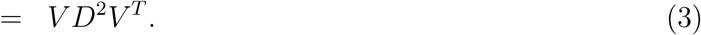

Setting 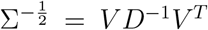, and evaluating the covariance between columns following the whitening transform gives identity covariance according to

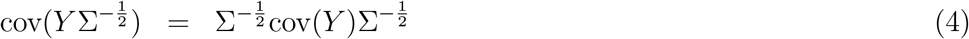

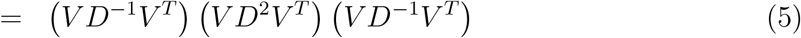

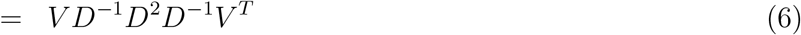

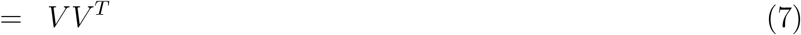

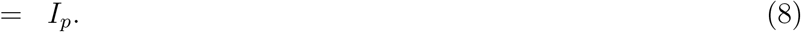

### 5.2 Evaluating whitening performance

Let *Â* be the whitening transformation learned from the observed data so that

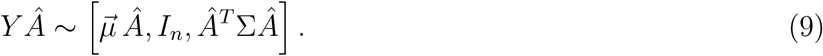

When *Â* ^*T*^ Σ *Â* = *I*_*p*_, as in Equation (2), this transformation produces data *Y Â* whose columns are *exactly* uncorrelated when *Â* is estimated from *Y*. Yet this condition can only be satisfied when the sample covariance matrix, 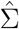 is invertible, and this requires *n > p*. In high dimensional data with *n < p*, so some form of regularization is needed. An exact whitening transformation no longer exists in these cases, so we want to learn *Â* from the data so that *Â* ^*T*^ Σ *Â* is ‘close’ to the identity matrix by some metric. Formally, we could seek a transform so that ∥ *Â* ^*T*^ Σ *Â* − *I*_*p*_ ∥_*F*_ is small, where ∥. ∥_*F*_ is the Frobenius norm of a matrix. This is closely related to the root mean squared error of the difference between the covariance of the transformed data and the identity matrix.

The Frobenius norm of a matrix *M*_*p×p*_ is defined as

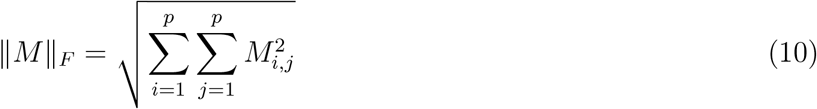

so that scaling by *p* gives

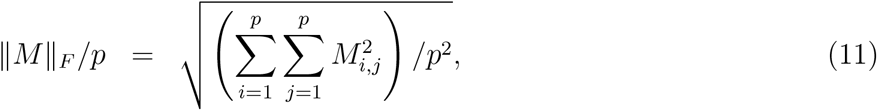

which is the root mean squared error when *M* = *Â* ^*T*^ Σ *Â* − *I*_*p*_.

### 5.3 Regularizing the correlation matrix and retaining sample variances

The approach described so far shrinks all diagonal variance entries towards a single value *ν*. While this is reasonable when the variances are all equal, most covariance matrices in practice have heterogeneous variances. Instead, we can factor out the variance terms, shrink the correlation matrix and then rescale by the variances. Let 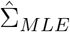 have variance 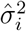 in the *i*^*th*^ entry on the diagonal, let *Z* be a diagonal matrix with 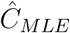 in the *i*^*th*^ entry and let *Ĉ*_*M LE*_ be the corresponding correlation matrix estimate so that the covariance can be decomposed according to 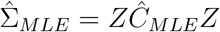. The regularized covariance matrix retaining the sample variances along the diagonal is then

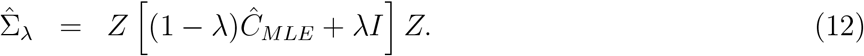

The approach described in the main text corresponds to setting *Z* = *I* and estimating *ν* from the data. Alternatively, a second approach corresponds to estimating the standard deviation matrix, *Z*, from the data and setting *ν* = 1. We omit the variance terms from the notation throughout for the sake of simplicity.

### 5.4 Estimation of shrinkage parameter

Setting the value of the shrinkage parameter *λ* is a key component for any convex combination method and many methods have been proposed under different, often nonparametric, assumptions [25, 26, 34, 35, 43, 62]. Following the probabilistic formulation, we take advantage of the Gaussian Inverse Wishart model and use an empirical Bayes approach to estimate the shrinkage parameter after marginalizing out the covariance. The probability of the observed data can be marginalized with respect to the unknown covariance term Σ according to

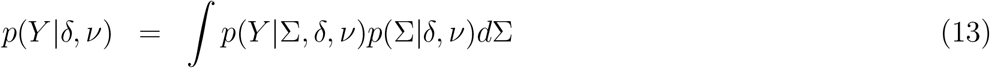

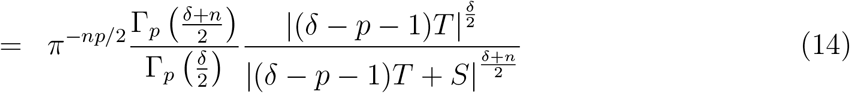

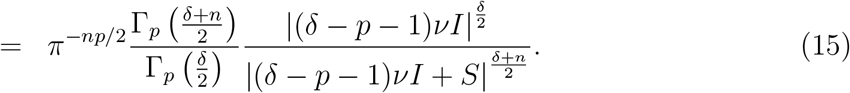

where *S* = *Y Y* ^*T*^, and we focus on the case where the target matrix is a scaled identity matrix so that *T* = *νI*.

The empirical Bayes estimates of *δ* and *ν* can be obtained from the observed data according to

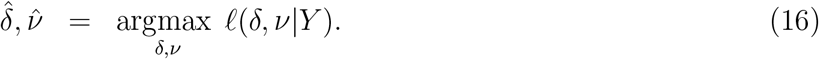

Where

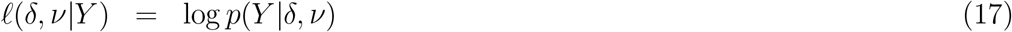

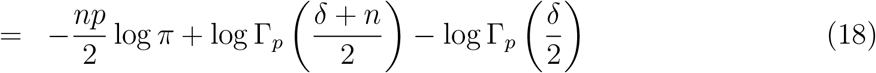

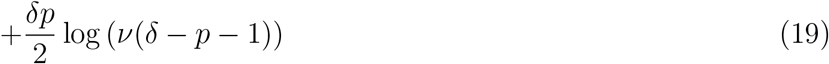

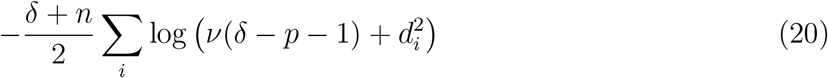

where 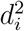 are the eigen values of *S*. Since 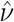 can be estimated in closed form from the data as the mean diagonals of *S*, and 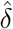 can be obtained subsequently using a line search, 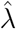 can then be obtained from 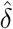. Importantly, this function has one parameter and the data enters only through the eigen-values *d*^2^ and the prior scale, *ν*.

#### 5.4.1 Low rank approximation

With the full spectral decomposition, the estimate of the correlation matrix

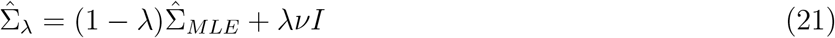

has diagonal values equal to 1. But when the data is approximated by a truncated singular value decomposition, 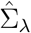 no longer has diagonals of value 1 when simply setting *ν* = 1. Instead, we set the value of *ν* so that the mean of the diagonals of 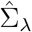 is closest to 1.

Let the target sum of the diagonal values of 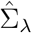 be *p*, and let

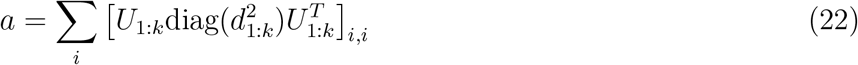

be the sum of the diagonals of the low rank approximation to the correlation matrix, 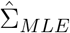. Therefore, the sum of the diagonals of 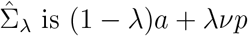.

Setting this value equal to the desired sum of the diagonals, *p*, and solving for *ν* gives

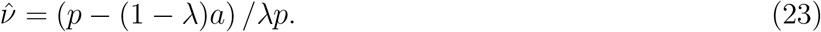

This value is estimated *within* each evaluation of *ℓ*(.) so that the function has just 1 parameter.

### 5.5 Properties as a function of *λ*

Here we can examine the behavior of the transformation as a function of *λ*. First, all *modified* singular values are positive, even in cases where the corresponding singular value of the observed data is zero. This is intuitive since all eigen-values of the implied covariance matrix 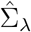 must be positive for it to be invertible.

Based on results above, setting *λ* = 0 corresponds to the algorithmically defined whitening transformation using the sample covariance matrix where there is no influence from the prior. Indeed, Equation (15) shows that this corresponds to setting all singular values of the transformed data to 1, thus scaling the sample variances in all dimensions to be equal. At the other extreme, setting *λ* = 1 divides each singular value by 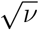 so the singular values of the transformed data are just scaled versions of the original values, and thus retains the same correlation structure. For intermediate values of *λ* that are used in practice, they are shrunk using *ν* as a ‘pivot’: values greater then *ν* are shrunk towards the pivot and values less than *ν* are inflated towards the pivot.

As *λ* → 1 the prior is increasingly concentrated around the target matrix and recovers the target matrix in the limit. As *λ* → 0, this corresponds to an infinitely diffuse prior and recovers the sample covariance matrix in the limit. As *n* increases for fixed *p*, a good estimator of *λ* should converge to 0 so the posterior expectation is an asymptotically consistent estimator of the covariance [33].

Mapping from the observed to the modified singular values depends on *λ*, which in turn depends on *n* and *p*, and 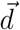. Small sample sizes require stronger regularization with larger *λ* values. In this case, the modified singular values retain some level of variation so the whitening transform is approximate. With increasing sample size, the effect of the prior decreases so that *λ* → 0 as *n* → ∞ with fixed *p*, so the modified singular values approach the observed singular values. At the same time,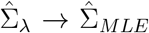 and 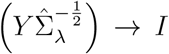 so that the whiting transform is asymptotically exact, and approximate for finite sample size.

The out-of-sample performance of the whitening transform has received less attention. For small sample size, the whiting transform estimated from the sample covariance can give very poor out-of-sample performance and can in fact increase the correlation between features. Regularization produces better estimates of the true singular values [63] and we demonstrate that it can also substantially improve out-of-sample whitening performance.

### 5.6 Interpretation of shrinkage parameter

The sample covariance 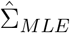 requires *p*(*p* − 1)*/*2 off-diagonal covariance parameters. Since the posterior expectation is a convex combination between prior and sample covariance, we can compute the effective number of parameters of the posterior estimate, 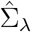, as a function of the *λ* shrinkage parameter.

Following Ye [49], the effective number of parameters is determined by the degree to which changes in the observed data change the estimator. Letting 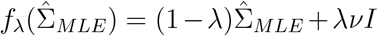 be the mapping from the sample covariance to the posterior covariance, Ye [49] defines the effective number of parameters as

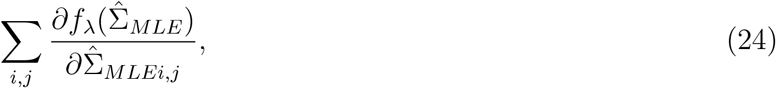

which is interpretable as the sum of the degree to which changes in the sample covariance change the posterior estimate. Considering only the off-diagonal covariance terms,

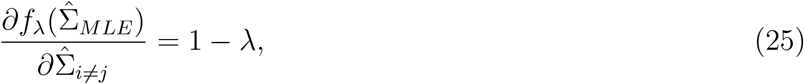

so each covariance term uses 1 − *λ* effective parameters. Therefore the effective number of parameters used to estimate the covariance terms is (1 − *λ*)*p*(*p* − 1)*/*2. Compared to the sample covariance, the effective number of parameters is thus reduced by a factor of 1 − *λ*.

As the number of features, *p*, increases with fixed sample size 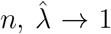 to apply stronger shrinkage [33]. While the effective number of parameters for the sample covariance matrix grows quadratically with *p*, it grows slower than quadratically for the shrinkage model since *λ* increases with *p* for fixed *n*.

### 5.7 Scalar summaries of the correlation or covariance matrix

The condition number of 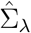 is defined as the ratio between the largest and smallest eigen-values:

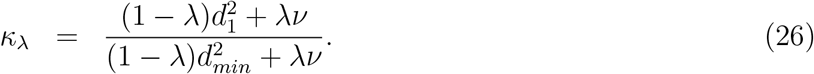

When the sample covariance is low rank, the smallest sample eigen-value is zero so the condition number has the simple form

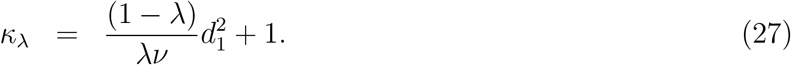

as long as *λ >* 0. Instead of regularizing the condition number using an ad hoc tuning parameter, or estimating it by cross validation, in the probabilistic model *λ* is a function of *n, p* and 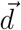. As *λ* increases, more weight is put on the target matrix, and the condition number approaches its minimum value of 1.

The effective variance is defined as the geometric mean of the eigen-values and satisfies a number of properties for a metric of multivariate variability [51]. This is defined as |Σ_*λ*_ |^1*/p*^, but can also be computed directly from the eigen-values. The effective number of features is analogous to the effective sample size based on the sum of all elements in the inverse correlation matrix [52] which is defined as 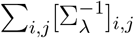. The average correlation can be computed by evaluating the sum of the off-diagonal entries of the correlation matrix divided by the number of entries. The effective number of features and average correlation can be computed efficiently using the implicit correlation matrix as described below. These values can be computed for the Σ_*MLE*_ matrix by setting *λ* = 0.

### 5.8 Scaling of standard whitening transform

Constructing 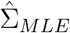 is *O* (*np*^2^), requires *O* (*p*^2^) memory, and evaluating its eigen decomposition is *O* (*p*^3^). Finally, explicitly creating the whitening matrix is *O* (*p*^2^), and evaluating the matrix product is *O* (*np*^2^). Estimating *λ* with the probabilistic model is *O* (*p*), but other methods look at the entire dataset so require at least *O* (*np*) time. Estimating *ν* is negligible in comparison, and applying shrinkage to the standard whiting transformation is also negligible. This allows us to consider the standard whitening transformation as a special case of the probabilistic model with *λ* = 0. The time complexity of the entire procedure is therefore *O* (*p*^3^ + *np*^2^). The standard algorithm becomes very expensive for *p >* 100 and can be intractable for *p >* 2000 features.

### 5.9 Accelerating transform using properties of linear algebra

#### Theorem 1

*Let C be a p* × *p symmetric positive semi-definite matrix, with k* ≤ *p positive eigen values. Let the eigen decoposition of C be U diag*(*s*)*U* ^*T*^ *where s*_*i*_ = 0 *for i > k. Let the eigen-vectors be partitioned as U* = [*U*_1_ *U*_2_], *and the eigen-values be partitioned as s* = [*s*_1_ *s*_2_] *to separate the nonzero and zero components. Let raising a matrix to an exponent α correspond to raising the eigen-values of the matrix to α. Then*

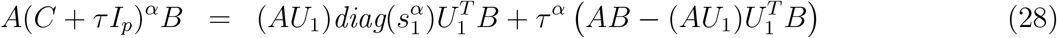

*where U*_1_ *and s*_1_ *indicate the first k eigen-vectors and values, respectively, and A*_*a×p*_ *and B*_*p×b*_ *are arbitrary matrices, τ is a positive scalar, and I*_*p*_ *is the p-dimensional identity matrix*.

*Proof*. The derivation follows from standard properties of linear algebra, the fact that the matrix of eigen-vectors, *U*, is orthonormal, and Lemma 1.1 below.

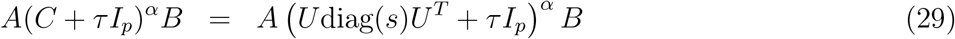

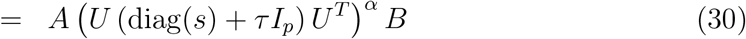

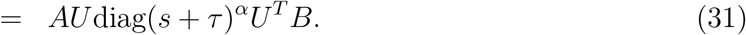

While this formula removes *U* from within the exponentiated term, it still includes *p* − *k* eigen-values that are zero along with their corresponding eigen-vectors. Partitioning the non-zero and zero terms of *s* and *U* and into separate components produces an expression using only non-zero terms.

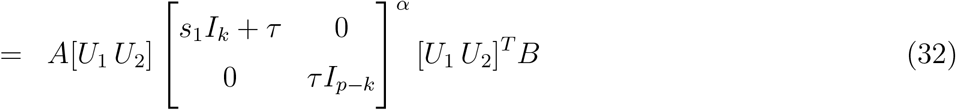

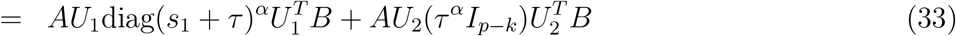

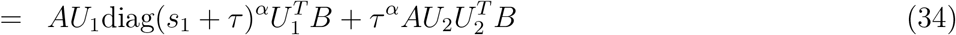

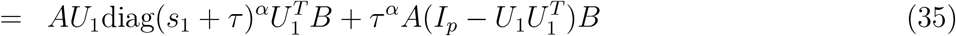

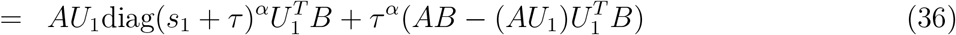

#### Corollary 1.1

*This allows efficient evaluation of A*(*C* +*τ I*_*p*_)^*α*^*B for many values of τ by precomputing s*_1_, *U*_1_, *AU*_1_ *and* 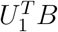.

#### Lemma 1.1

*Since U is orthonormal*,

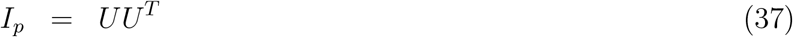

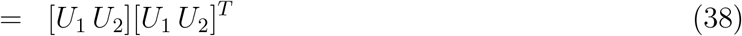

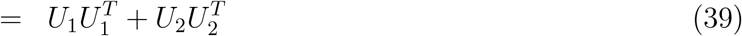

*Rearranging, it follows that* 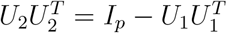.

### 5.10 Application to imputing z-statistics

Consider a matrix *X*_*n×p*_ of *p* genetic variants from *n* individuals, and a vector of continuous trait values *y*. Standard analysis in genome-wide association studies performs a linear regression to test association of each genetic variant with the trait. Ignoring covariates here for simplicity, this estimates a regression coefficient for the *j*^*th*^ variant as 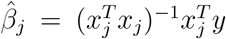, and produces a corresponding z-statistic, *z*_*j*_. Within a local genomic region, genetic variants can be highly correlated due to linkage disequilibrium (LD). Therefore, even if the *k*^*th*^ genetic variant is not directly observed, its z-statistic can be imputed from the set of observed z-statistics and the correlation structure between variants in the region. Following Pasaniuc, et al. [16], let Σ_*p×p*_ be the correlation matrix between all pairs of genetic variants, and partition the genetic variants into two sets with *j* storing indices with observed z-statistics and *k* storing indices without z-statistics. We would like to impute the values of *z*_*k*_ given observed values of *z*_*j*_ and Σ. If we model the z-statistics as a multivariate Gaussian

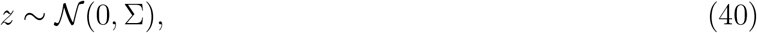

then the conditional distribution of *z*_*k*_ given *z*_*j*_ is

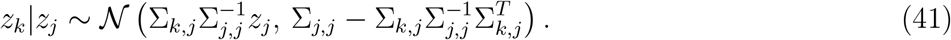

The imputed z-statistics is then

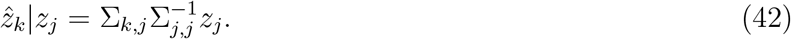

This also works for logistic regression in the case / control studies.

So far, we have assumed that the correlation matrix Σ is known. But in practice, this ‘LD matrix’ is estimated from a ‘reference panel’ of a modest number of individuals that are not included in the genome-wide association study. Therefore, inverting Σ_*j,j*_ requires some regularization as *p* becomes large in comparison to *n*.

In the context of the current work, Equation (42) can be in interpreted as an out-of-sample whitening transformation of the observed z-statistics, *z*_*j*_, and the cross-correlation matrix, Σ_*k,j*_. Letting 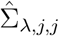 be the regularized estimate of the correlation matrix, then the imputed z-statistic is

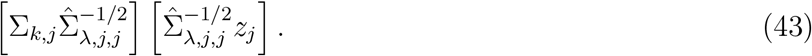

Applying this transformation by producing 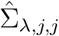 directly is cubic time in *p*_*j*_, the number of genetic variants in set *j*. But applying the high dimensional whitening transformation of Equation (19) reduces this time substantially to *O* (*n*^2^*p*_*j*_) when *n < p*_*j*_. Imputation then depends on the observed z-statistics the cross-correlation matrix, and the SVD of *X*.

### 5.11 Imputing z-statistics for tandem repeats

The VCF files from the EnsembleTR v4 [57] reference panel including donors from 1000 Genomics Project was obtained from https://github.com/gymrek-lab/EnsembleTR. Summary statistics from a study of genetic regulation of gene expression in the human brain [58] was downloaded from https://www.synapse.org/Synapse:syn25592272, and z-statistics from the fixed effects meta-analysis were used here.

Due to the complexity of harmonizing tandem repeats identified in sequence data, the EnsembleTR reference panel [57] often has multiple alternatively alleles at a site. For example, consider the hexanucleotide repeat near C9orf72 which is encode with 25 alternative alleles (**Table S1**). When multiple alleles are observed at a site, the alleles are modified to have multiple biallelic variants at the same site using the command: `bcftools norm -m - ${VCF}`. The z-statistics are imputed for each biallelic variant, and then combined into a single z-statistic for that site using a random effects metaanalysis that models the correlation structure between the variants [64]. This is implemented using the `RE2C()` function in the `remaCor` package available at https://cran.r-project.org/package=remaCor.

## 6 Supplementary Tables

**Table 1.**
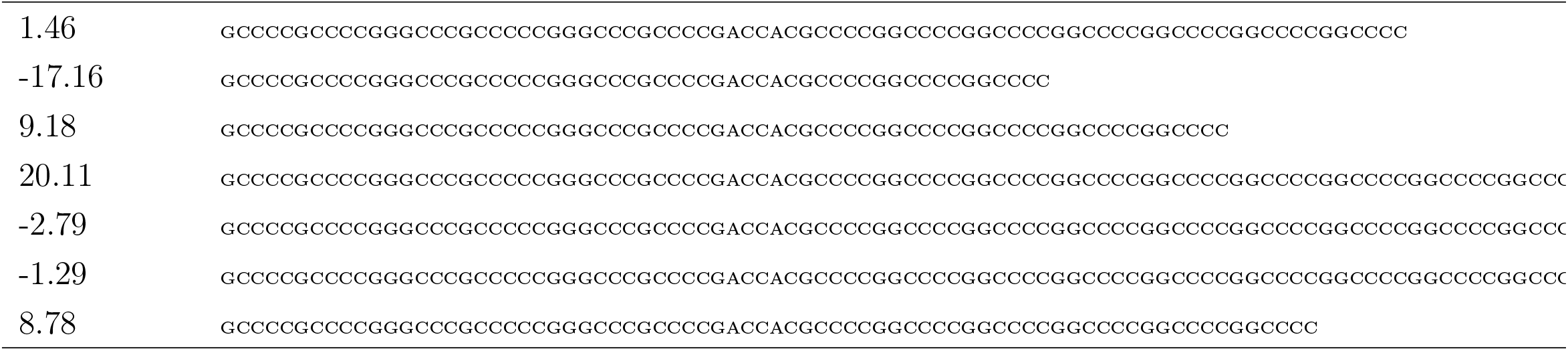
Imputed z-statistics for tandem repeat EnsTR:chr9:27573485 associated with expression of C9orf72. Due to the complexity of harmonizing tandem repeats identified in sequence data, the EnsembleTR reference panel [57] encodes the known hexanucleitude repeat near C9orf72 as a 62bp reference allele (GCCCCGCCCCGGGCCCGCCCCCGGGCCCGCCCCGACCACGCCCCGGCCCCGGCCCCGGCCCC) with 25 alternative alleles. The 7 most common are shown here with their imputed z-statistics.

## 7 Supplementary Figures

**Figure S1.**
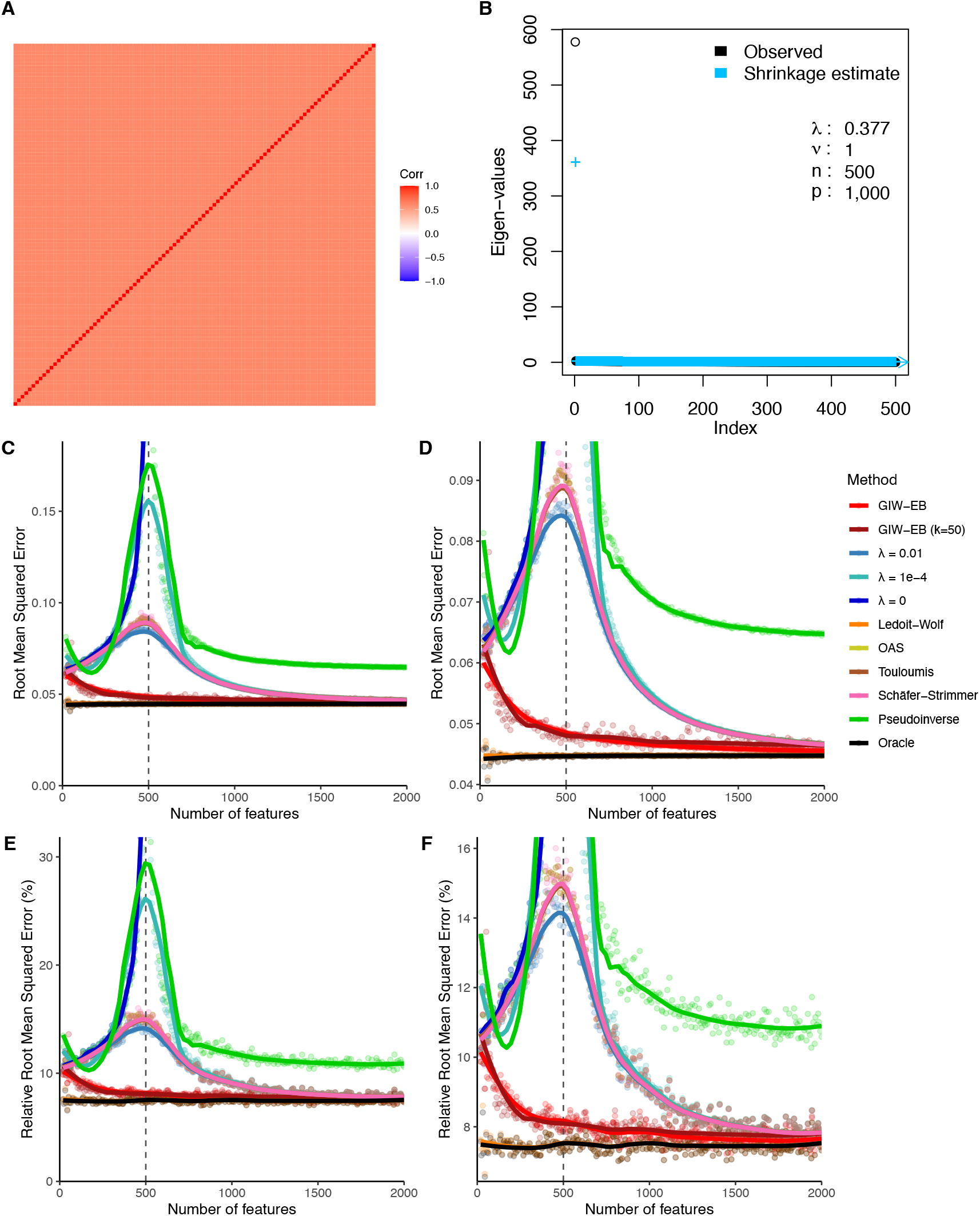
Out-of-sample performance of whitening transformation. Scenario 1: constant correlation. Details are given in **Figure 3. A**) Constant correlation matrix with *ρ* = 0.6 used in this simulation. **B**) Observed and shrunk eigen-values from simulation of *p* = 1000 features. **C**,**D**) rMSE of the whitening transformation for an increasing number of features with a zoom-in on the y-axis. **E**,**F**) rMSE scaled by the rMSE of the untransformed data with a zoom-in on the y-axis. Points indicate observed values and lines show loess smooth.

**Figure S2.**
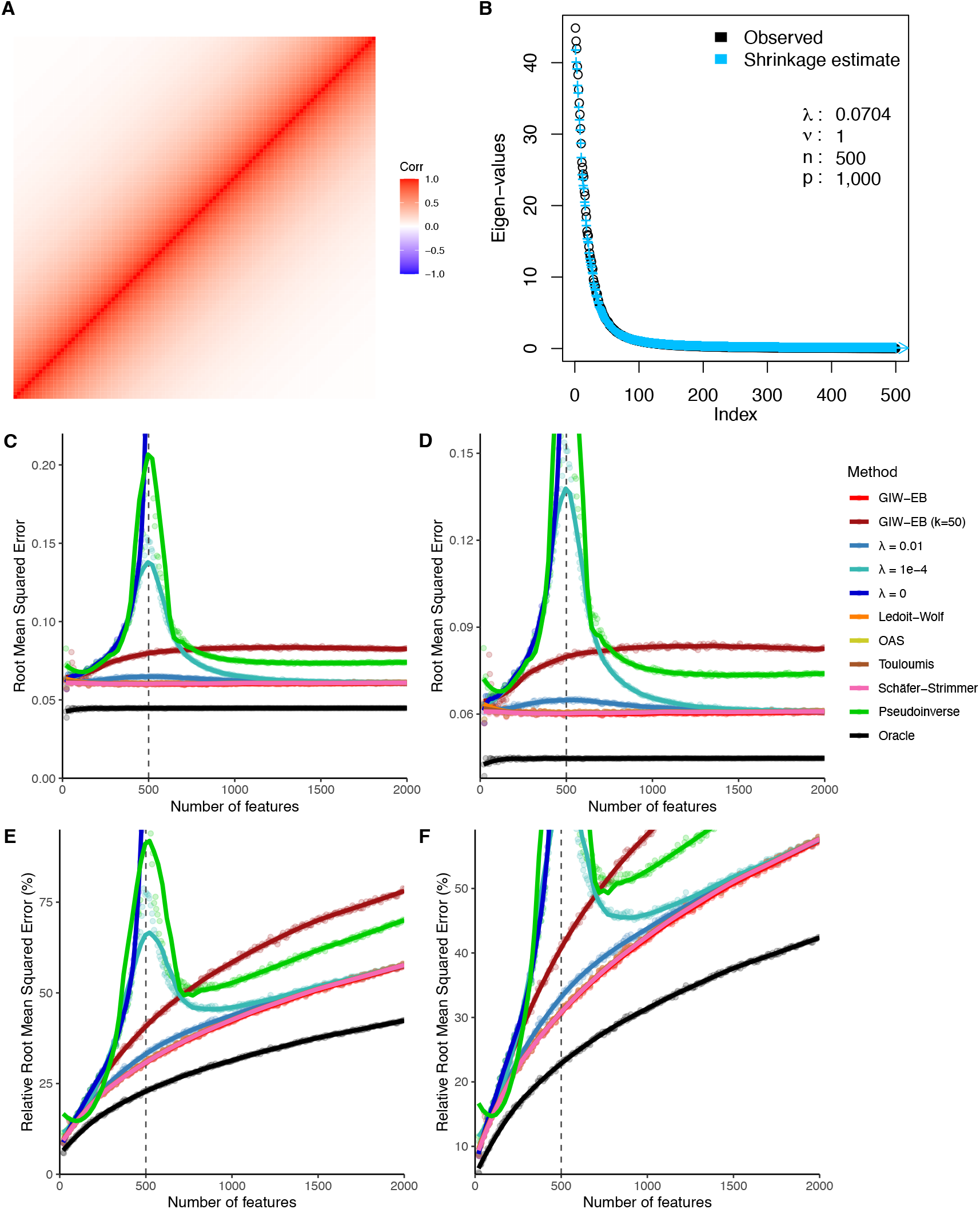
Out-of-sample performance of whitening transformation. Scenario 2: auto-correlation. Details are given in **Figure 3. A**) Auto-correlation matrix with *ρ* = 0.95 used in this simulation. **B**) Observed and shrunk eigen-values from simulation of *p* = 1000 features. **C**,**D**) rMSE of the whitening transformation for an increasing number of features with a zoom-in on the y-axis. **E**,**F**) rMSE scaled by the rMSE of the untransformed data with a zoom-in on the y-axis. Points indicate observed values and lines show loess smooth.

**Figure S3.**
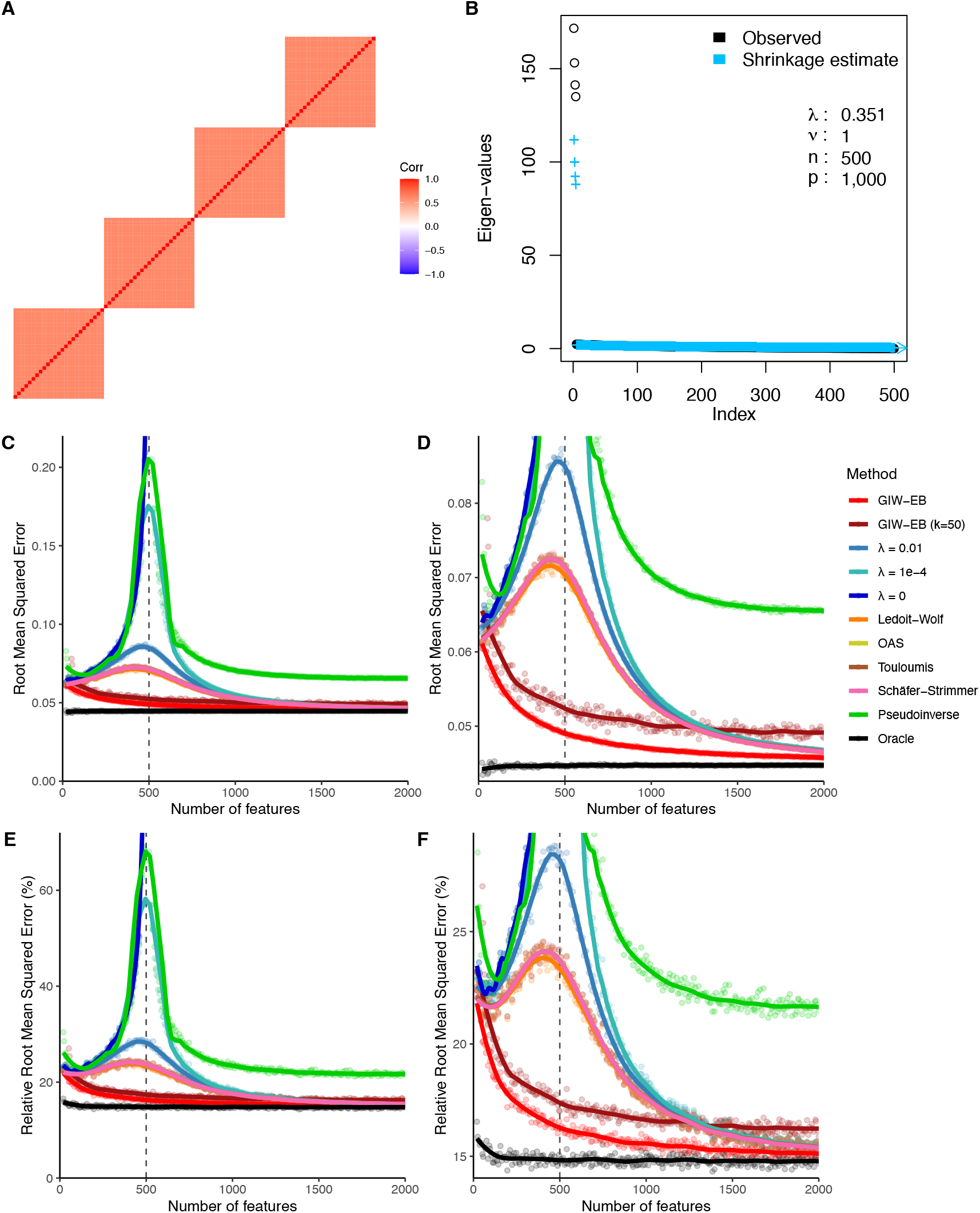
Out-of-sample performance of whitening transformation. Scenario 3: block constant correlation. Details are given in **Figure 3. A**) Block constant correlation matrix with *ρ* = 0.6 used in this simulation. **B**) Observed and shrunk eigen-values from simulation of *p* = 1000 features. **C**,**D**) rMSE of the whitening transformation for an increasing number of features with a zoom-in on the y-axis. **E**,**F**) rMSE scaled by the rMSE of the untransformed data with a zoom-in on the y-axis. Points indicate observed values and lines show loess smooth.

**Figure S4.**
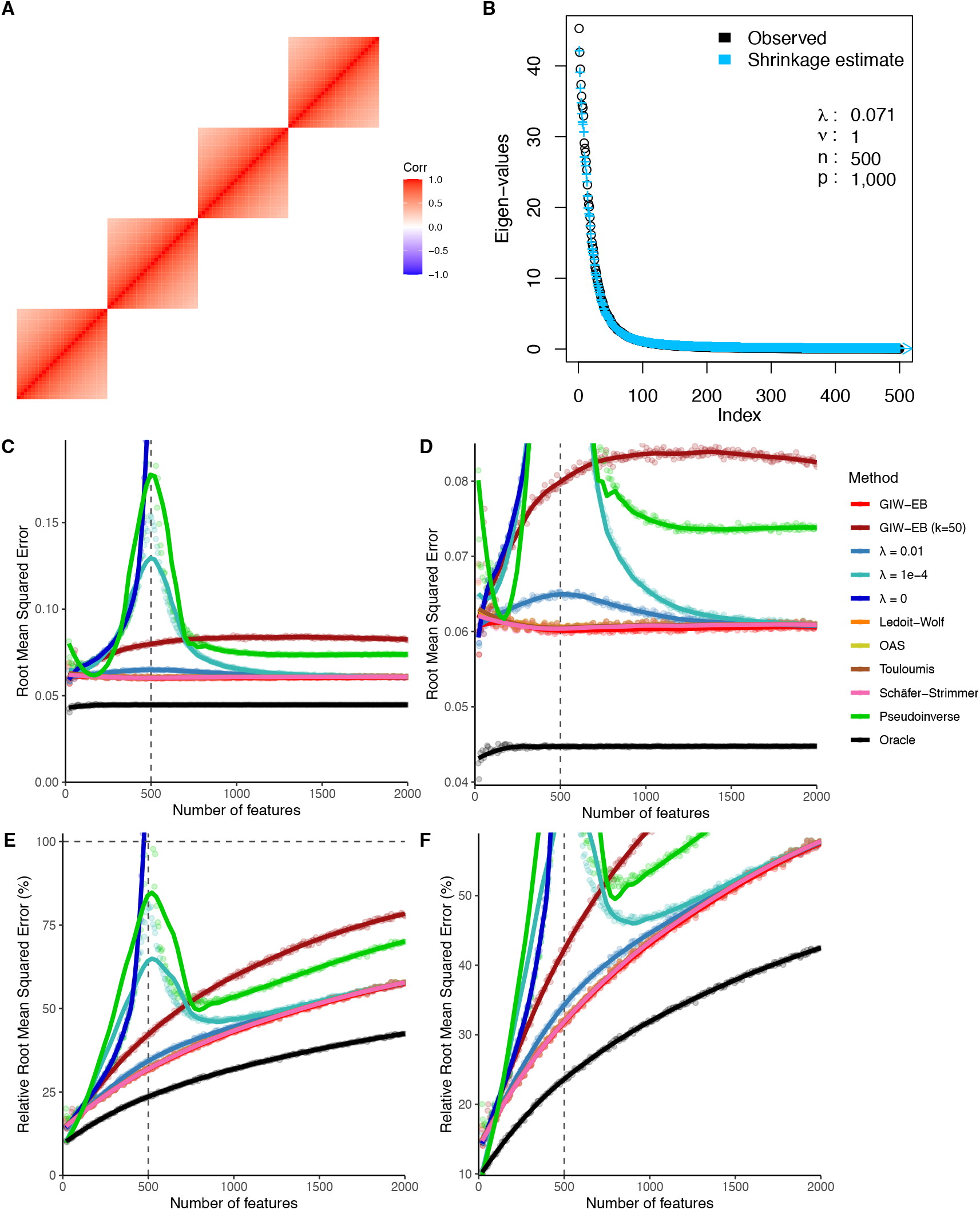
Out-of-sample performance of whitening transformation. Scenario 4: block auto-correlation. Details are given in **Figure 3. A**) Block auto-correlation matrix with *ρ* = 0.95 used in this simulation. **B**) Observed and shrunk eigen-values from simulation of *p* = 1000 features. **C**,**D**) rMSE of the whitening transformation for an increasing number of features with a zoom-in on the y-axis. **E**,**F**) rMSE scaled by the rMSE of the untransformed data with a zoom-in on the y-axis. Points indicate observed values and lines show loess smooth.

**Figure S5.**
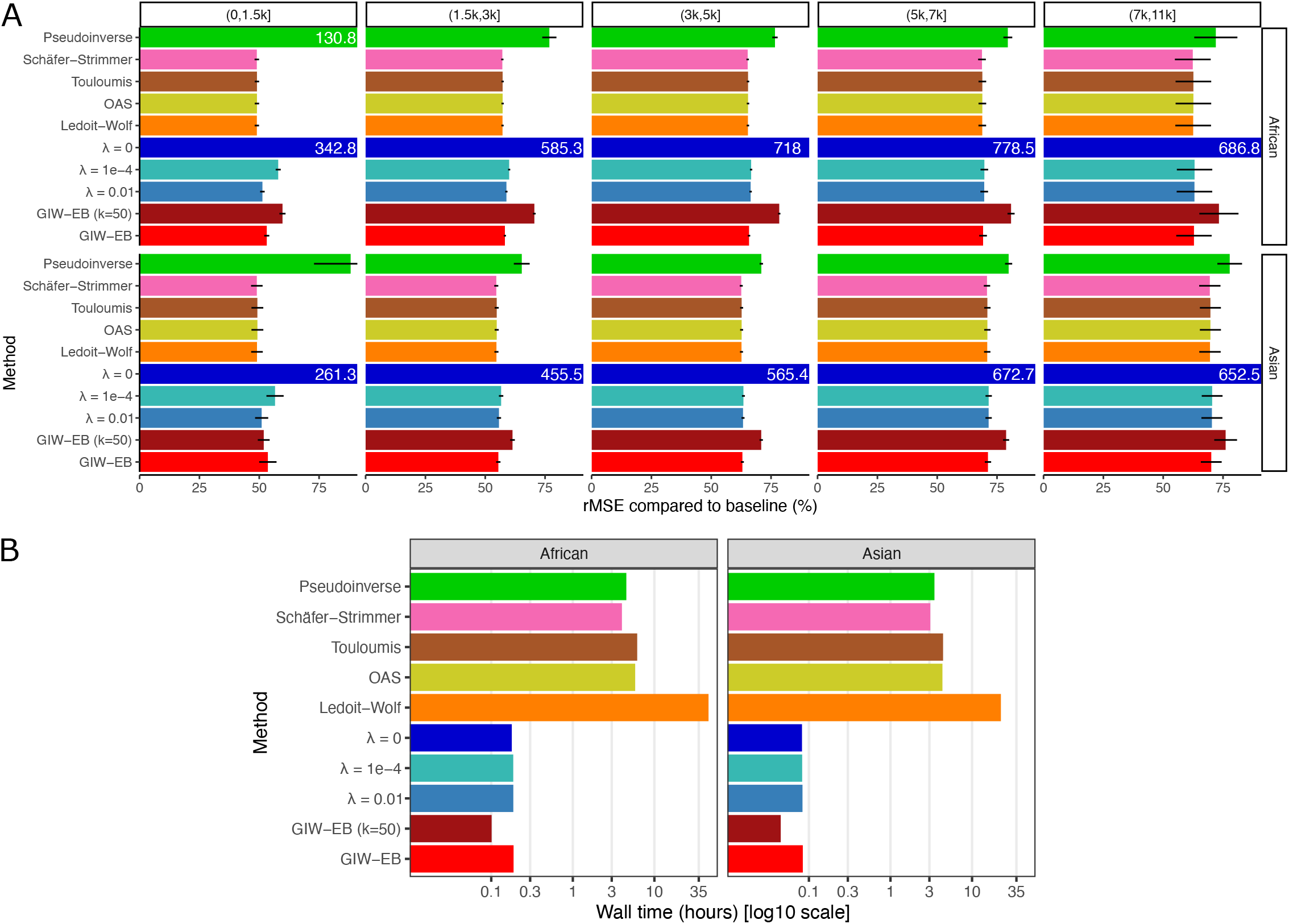
Out-of-sample performance of whitening transformation on genotype data from the 1000 Genomes Project. **A**)Root mean squared error (rMSE) compared to baseline for African and Asian individuals stratified by the number of genetic variants in the LD window. Values indicate the values averaged across windows, and bars indicate the 95% confidence interval. When normalized rMSE values exceed the axes, the value is shown in white text. **B**) Run time of each method shown on a log_10_ scale.

## Footnotes

1 We use the matrix normal here in order to model covariance between columns while having independent rows, but the results remain unchanged when transposing the observed data and using the multivariate normal notation instead.

